# In vitro evaluation of probiotic properties of lactic acid bacteria isolated from some traditionally fermented Ethiopian food products

**DOI:** 10.1101/574194

**Authors:** Guesh Mulaw, Tesfaye Sisay, Diriba Muleta, Anteneh Tesfaye

## Abstract

Probiotics are live microorganisms which when consumed in large number together with a food promote the health of the consumer. The aim of this study was to evaluate *in vitro* probiotic properties of lactic acid bacteria (LAB) isolated from traditional Ethiopian fermented *Teff injera* dough, *Ergo* and *Kocho* products. A total of 90 LAB were isolated, of which 4 (4.44%) isolates showed 45.35-97.11% and 38.40-90.49% survival rate at pH values (2, 2.5 and 3) for 3 and 6 h in that order. The four acid tolerant isolates were found tolerant to 0.3% bile salt for 24 h with 91.37 to 97.22% rate of survival. The acid-and-bile salt tolerant LAB isolates were found inhibiting some foodborne test pathogenic bacteria to varying degrees. All acid-and-bile tolerant isolates displayed varying sensitivity to different antibiotics. The *in vitro* adherence to stainless steel plates of the 4 screened probiotic LAB isolates were ranged from 32.75 to 36.30% adhesion rate. The four efficient probiotic LAB isolates that belonged to *Lactobacillus* species were identified to strain level using 16S rDNA gene sequence comparisons and namely were *Lactobacillus plantarum* strain CIP 103151, *Lactobacillus paracasei* subsp. tolerans strain NBRC 15906, *Lactobacillus paracasei* strain NBRC 15889 and *Lactobacillus plantarum* strain JCM 1149. The four *Lactobacillus* strains were found to have potentially useful to produce probiotic products.

## Introduction

Worldwide a variety of fermented food products are produced, which contribute significantly to the diets of many people (1). Fermented food products are used to describe a special class of food products characterized by various kinds of carbohydrate breakdown in the presence of probiotic microorganisms; but seldom is carbohydrate the only constituent acted upon (2). Fermented food and beverage products have emerged as not only the source of nutrition but also as functional and probiotic foods, which besides nutritional value have health effects or provide protection against foodborne diseases.

The problem of foodborne diseases (FBD) is multi factorial and their prevention and control require a multidisciplinary approaches that involve human beneficial live microbes (probiotics) in order to combat these pathogens and their associated health risks (3). Several *in vitro* studies indicated that the growth of foodborne pathogenic microbes is inhibited by probiotic lactic acid bacteria (4-6). The consumption of large number of probiotics live microorganisms together with a food fundamentally promote the health of the consumers (7). Accordingly, lactic acid bacteria (LAB) are Generally Recognized as Safe (GRAS) by WHO and play an important role in the process of fermentation of food by inhibiting spoilage/pathogenic bacteria and by producing excellent flavor, aroma, and texture of fermented foods (8,9). LAB are candidate probiotic bacteria that are widely distributed in nature and can be used in the food industry (10).

LAB could be isolated from many kinds of sources such as milk products, fermented foods, animal intestines or freshwater fishes, soil samples, sugar cane plants, and poultry farms (11). The most common types of probiotic LAB include different *Lactobacillus* spp. (*Lb. acidophilus, Lb. johnsonii, Lb. casei, Lb. rhamnosus, Lb. gasseri* and *Lb. reuteri*) and genus *Bifidobacteria* (*Bf. bifidum, Bf. lactis, Bf. Longum* and *Bf. infantis*) (Arthure *et al*., 2002; Chassard *et al*., 2011). LAB are also useful in the treatment of various diseases caused by drug resistant pathogenic microbes (12). Probiotic microbes provide nutrients, enhance growth, produce enzymes, inhibit pathogens and enhance immune responses (13).

In Ethiopia, traditionally fermented food products are prepared at household level and the consumption of these fermented food products is commonly practiced. Therefore, some studies on isolation and screening of antibacterial producing lactic acid bacteria from traditionally fermented foods were undertaken by many workers (14,15). Likewise, Tesfaye, Mehari (4) have revealed that the antagonistic effect of lactic acid bacterial strains either as pure or defined mixed-cultures, against some foodborne pathogens during fermentation and storage of fermented milk. However, there are still few research data available on characterization of probiotic LAB. Most of the traditionally fermented products of Ethiopia are consumed without further heat processing which can be considered as ideal vehicles to carry probiotic bacteria into the human gastrointestinal tract.

Probiotic strains isolated from traditionally fermented foods and drinks could have application as a starter cultures for a large scale production of the traditional product and have a desirable functional property for their application as probiotics against foodborne pathogens. Therefore, this study attempts to evaluate the *in-vitro* probiotic properties of LAB isolated from three traditionally fermented Ethiopian food products such as *Teff dough, Ergo* and *Kocho* with respect to their probiotic properties against some food-borne pathogens.

## Materials and methods

### Sample collection

Traditionally fermented food products (*Teff dough, Kocho* and *Ergo*) were obtained from Addis Ababa and its surroundings, Ethiopia. Each sample (200 g) was aseptically collected by using sterilized containers. The samples were brought to the laboratory with ice box and stored in a refrigerator at +4°C until further analysis was carried out. *Ergo* is locally fermented milk product. *Teff dough* is made by fermenting teff (*Eragrostis tef*) flour which is used to prepare thin pancake like product with many eyes known as *injera*. *Kocho* is product which is prepared from decorticated and pounded pulp of enset plant (*Enset ventricosum*), which is further mixed and kneaded into a mash and fermented in a pit.

### Isolation and purification of LAB from traditional fermented foods

For isolation of LAB, 25 ml or 25 g of each sample of traditionally fermented foods (*Teff dough, Kocho* and *Ergo*) was mixed with 225 ml of separate sterile peptone-water (0.1% W/V). Then, a sequential decimal dilution of the homogenate was obtained. From the appropriate dilutions, 0.1 ml aliquots were spread plated on duplicate pre-dried surfaces of MRS (de-Mann, Rogosa and Sharp) agar (Oxoid) plates. The inoculated plates were incubated under anaerobic condition using anaerobic jar (BBL, GasPak Anaerobic Systems) at 37°C for 48 hours. Then, 10-20 distinct colonies were randomly picked from countable MRS plates for further purification. The isolated colonies of LAB were transferred into about 5 ml MRS broth (Oxoid) and purified by repeated streaking on MRS agar. Pure cultures of LAB were then streaked onto slants of MRS agar, and stored at +4°C for further characterization (16).

### Designation of the LAB isolates

The LAB isolates were designated with E for *Ergo*, T for *Teff dough* and K for *Kocho*, followed with different numbers.

### Confirmation tests of LAB isolates

#### KOH test

KOH test was used to determine gram reaction of LAB isolates. LAB cultures were grown on MRS agar at 37°C for 24 h under anaerobic conditions. A drop of 3% aqueous KOH was placed on a clean slide. Using a sterile loop, visible cells from fresh cultures were transferred to the drop of 3% KOH. The cells and KOH were mixed thoroughly on the slide and stirred constantly over an area about 1-2 cm^2^. The isolates, which did not give viscid, were selected since, lactic acid bacteria (LAB) are known as Gram positive cells (17).

#### Catalase test

Overnight cultures of isolates were grown on MRS agar at +37°C for 24 h under anaerobic conditions. Catalase test was conducted by dripping two drops of hydrogen peroxide (3%) on 24 h old cultures on a glass slide. Catalase test positive reaction characterized by the formation of oxygen bubbles that indicate the production of catalase enzyme by the test bacterium. Therefore, the isolates, which did not give gas bubbles were selected for subsequent activities.

#### Spore staining

Gram-positive and catalase-negative isolates were grown on MRS agar at +37°C for 24 h under anaerobic conditions. Spore staining procedure was applied (18). After, spore staining technique, the endospore formulation was examined under light microscopy using oil immersion objectives. The isolates which did not form endospores were selected for further analysis.

#### In *vitro* characterization of probiotic properties

For the determination of probiotic properties (tolerance to low pH, tolerance against bile salt, antibiotic susceptibility, the antimicrobial activity and bacterial adherence to stain steel plates) of the isolates were carried out.

#### Tolerance to low pH

The isolates were grown separately overnight in 5 ml MRS broth at +37°C under anaerobic conditions. A volume of 1ml of log 7 cfu/ml of each overnight grown culture was inoculated in to 10 ml of MRS broth to give initial inoculum level of log 6 cfu/ml. The culture was then centrifuged at 5000 rpm for 10 min at +4°C. The pellets were washed twice in phosphate-saline buffer (PBS at pH 7.2). The pellets were re-suspended in 5 ml sterile MRS broth which was adjusted to pH values of 2.0, 2.5, and 3.0 using 1 N HCL to simulate the gastric environment. The test tubes were incubated for 3 and 6 hours at 37°C. After appropriate incubation period, 1ml of the culture was diluted in sterile 9 ml phosphate buffer (Sigma, St. Louis, USA) prepared according to the manufacturer’s instruction (0.1 M, pH 6.2) in order to neutralize the medium acidity. Briefly, a 100-µl aliquot of the culture and its 10-fold serial dilutions were plated on MRS agar medium. The inoculated plates were incubated at 37°C for 24 to 48 h under anaerobic condition using anaerobic jar (BBL, Gas Pack System). The grown LAB colonies were expressed as colony forming units per milliliter (cfu/ml). A positive control consisting of regular MRS broth inoculated with the culture was used (19). The survival rate was calculated as the percentage of LAB colonies grown on MRS agar compared to the initial bacterial concentration.

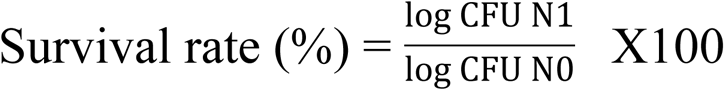

Where, N1: Viable count of isolates after incubation; N0: Initial viable count

#### Tolerance to bile salts

To estimate bile tolerance of Acid-tolerant LAB (those only were grown in pH 2.0, 2.5 and/or 3.0), the isolates were separately grown overnight in MRS broth at 37°C under anaerobic conditions (19). Each culture with the initial concentration of 10^6^ cfu/ml was then centrifuged at 5000 rpm for 10 min at 4°C. The pellets were washed twice in phosphate-saline buffer (PBS at pH 7.2). Cell pellets were re-suspended in sterile MRS broth supplemented with 0.3% (w/v) bile salt (Oxgall, USA). Samples were taken at 24 h from the onset of incubation to determine the survivability of cells as described previously (19). A positive control consisting of plain MRS broth without bile salts inoculated with each separate culture was simultaneously set up. After appropriate incubation, 1ml of each separate culture was diluted separately in sterile 9 ml phosphate buffer (Sigma, St. Louis, USA) prepared according to the manufacturer’s instruction (0.1 M, pH 6.2) in order to neutralize the medium. Concisely, a 100-µl aliquot of the culture and its 10-fold serial dilutions were plated on MRS agar medium. Plates were incubated at 37°C for 24 to 48 h under anaerobic condition using anaerobic jar (BBL, Gas Pack System). LAB counts were expressed in colony forming units per milliliter (cfu/ml). The survival rate was calculated as the percentage of LAB colonies grown on MRS agar compared to the initial bacterial concentration.

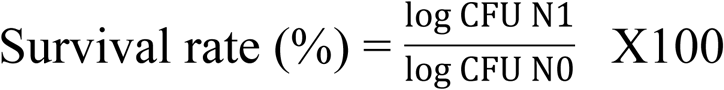

Where, N1: Viable count of isolates after incubation; N0: Initial viable count

#### Antimicrobial activity

Antibacterial activity of the acid bile-tolerant lactic acid bacteria strains against some foodborne pathogens was determined using the agar well diffusion method with some modifications the protocol indicated in Fontana, Cocconcelli (20). The test organisms (*Staphylococcus aureus, Listeria monocytogenes, Salmonella typhi* and *Escherichia coli)* were obtained from Ethiopian Public Health Institute (EPHI), Addis Ababa, Ethiopia.

The selected acid-bile-tolerant LAB isolates were inoculated from slants to fresh MRS broth containing 1% glucose and incubated overnight at 37°C. The overnight active culture broth of each isolate was centrifuged separately at 5000 rpm for 10 min at 4°C. The cell free supernatant from each separate culture was collected as crude extract for the antagonistic study against selected food borne pathogens. The pure cultures of foodborne pathogens were inoculated from slants to brain heart infusion broth. After 24 hour incubation at 37°C, a volume of 100 µl of inoculum of each indicator bacteria was swabbed evenly over the surface of nutrient agar plates with a sterile cotton swab. The plates were allowed to dry and a sterile cork borer (diameter 5 mm) was used to cut uniform wells in the agar. Each well was filled with 100 µl culture free filtrate obtained from each of the acid-bile tolerant LAB isolates. After incubation at 37°C for 24 to 48 hours, the plates were observed for a zone of inhibition (ZOI) around the well. The diameter of the inhibition zone was measured by calipers in millimeters and clear zone of 1 mm or more was considered positive inhibition (21,22). The experiment was carried out in triplicates and the activity was reported as diameter of ZOI ± SD.

#### Antibiotic susceptibility tests

Each of acid-bile-tolerant and antagonistic lactic acid bacteria isolate was assessed for its antibiotic resistance by disc diffusion method as described by Zhang, Wang (23) against some antibiotics that included ampicillin (10 μg/ml), erythromycin (15 μg/ml), streptomycin (10 μg/ml), kanamycin (25 μg/ml), and tetracycline (30 μg/ml). Thus, a volume of 100 µl of actively growing cultures of each acid-bile-tolerant and antagonistic lactic acid bacteria was swabbed evenly over the surface of nutrient agar plates with a sterile cotton swab. After drying, the antibiotic discs were placed on the solidified agar surface, and the plates were left aside for 30 min at 4°C for diffusion of antibiotics and then anaerobically incubated at 37°C for 24 to 48 h. Resistance was defined according to the disc diffusion method by using the above antibiotic discs and the diameters of inhibition zones were measured using calipers (24) and zone of inhibition (diameter in mm) for each antibiotic was measured and expressed as susceptible, S (≥21 mm); intermediate, I (16-20 mm) and resistance R (≤15.

#### Bacterial adhesion to stainless steel plates

The adherence assay of acid-bile-tolerant, antagonistic and antibiotic-sensitive lactic acid bacterial isolates was determined on stainless steel plates with some modifications of the protocol given in El-Jeni, El Bour (25). Briefly, LAB was cultured in sterile MRS broth. Thereafter, the overnight bacterial culture (500 µl) was deposited in a test tube, which was then filled with 450 µl of MRS broth, wherein the sterile stainless steel plate was deposited, and the test tubes were then incubated for 24 h at 37°C. The stainless steel plate was removed under aseptic conditions, washed with 10 ml of sterile 1% peptone water, and left for 5 min in a sterile 1% peptone water tube. The plate was then washed again in the same conditions and vortexed for 3 min in a sterile 1% peptone water tube (6 ml) consecutively to detach the bacterial cells adhering to the steel plate surface. Cell number was determined by counting on MRS agar after 24 h of incubation at 37°C. Simultaneously, the total initial cell numbers were estimated to calculate the percentage of adhered bacterial cells for each LAB.

#### Morphological, biochemical and physiological tests

The probiotic LAB isolates were identified according to their morphological, physiological and biochemical characteristics based on Bergey’s Manual (26).

#### Cell morphology

Overnight cultures were wet mounted on microscopic slide and examined under light microscope using oil immersion objectives. Cellular morphological criteria considered during examination were cell shape and cell arrangements.

#### Growth at different temperatures

Each of the overnight LAB culture of 50 µl was transferred into four separate tubes that contained 5 ml medium (modified MRS broth) containing bromecresol purple indicator at a concentration of 0.12g/l. After inoculation, two of the inoculated test tubes were incubated for 7 days either at 15°C and the other two test tubes at 45°C. During this incubation time, growth at any temperature was observed by the change of the growth medium (cultures) from purple to yellow.

#### Growth at different NaCl concentrations

LAB isolates were tested for their tolerance to different NaCl concentrations. For this purpose, 4% and 6.5% NaCl concentrations were used for testing. Similarly, test tubes with 5 ml of modified MRS broths containing bromecresol purple indicator was prepared according to the appropriate concentrations. Four test tubes with 4% and the other four test tubes with 6.5% NaCl were inoculated separately with 50 µl of 1% of each overnight culture of LAB and incubated at +37 °C for 7 days. The change of the color from purple to yellow was considered as a proof of the cell growth.

#### Arginine hydrolysis test

Arginine containing MRS medium and Nessler’s reagent (HgI4K2), a 0.09 mol/L solution of potassium tetraiodomercurate (II) (K2 [HgI4]) in 2.5 mol/L potassium hydroxide were used in order to test ammonia production from arginine. MRS containing 0.3% L-arginine hydrocloride was transferred into duplicate tubes with 5 ml amount and inoculated with each of 1% overnight cultures of LAB. Thereafter, tubes were incubated at +37°C for 24 h. A 100 μl of cultures were transferred onto a white background. Thereafter, the same amount of Nessler’s reagent was pipetted to the cultures. The change in color to bright orange indicates a positive reaction, while the yellow color indicates the negative reaction. A negative control, which does not contain arginine, was used as negative control.

#### Gas production from glucose

In order to determine the homonfermentative and heterofermentative characteristics of LAB isolates, CO_2_ production from glucose was determined in modified MRS broth containing inverted Durham tubes with 1% glucose. MRS broth (8 ml) in separate tubes containing 1% glucose with inverted Durham tubes were prepared and inoculated separately with 50 µl of 1% overnight fresh each LAB culture. Then the test tubes were incubated at +37 °C for 5 days. The presence of gas in Durham tubes during 5 days observation indicates CO_2_ production from glucose.

### Identification of probiotic LAB isolates using 16S rRNA Gene sequencing

#### Genomic DNA extraction

Genomic DNA was extracted from pure cultures (n=4) of potential probiotic LAB. Separately 1 ml of each pure liquid culture was centrifuged for 3 min at 10000 rpm. The supernatant was removed and the cells were re-suspended in 300 μl buffer (10 mM Tris-HCl, pH 8.0; 50 mM glucose, 10 mM EDTA). To the suspension, 3 μl lysozyme (10 mg/ml) was added and cells were lysed at 37°C for 60 min under occasional stirring of the tube content by overturning it. Lysing buffer (20 mM Tris-HCl, pH 8.0; 75 mM NaCl; 1% SDS; 10 mM EDTA) of 300 μl and 3 μl RNAse (10 mg/ml) were added to the mixture. The mixture was incubated at 37°C for 30 min, and then cooled on ice for 1 min. Then, 100 μl solution ammonium acetate (7.5 M) was added to the mixture and mixed on a vortex for 20 seconds was centrifuged at 13000 rpm for 5 min. The supernatant was transferred into clean 1.5 ml tubes and 300 μl isopropanol was added. Thereafter, the mixture was mixed by overturning for 1 min and stored at–20°C for 30 min. The mixture was centrifuged at 13000 rpm for 5 min. The supernatant was accurately decanted and the tubes were placed overturned on a clean filter. Then, 400 μl of 70% ethanol was added and mixed several times by overturning to wash the DNA sediment. Finally, the sediment was dried at 37°C for 15 min till ethanol drops disappeared completely. The dried sediment was dissolved in 30 μl TE buffer.

#### PCR amplification of 16S rDNA

For the amplification of 16S rDNA gene, the specific primers AMP_F 5’-GAG AGT TTG ATY CTG GCT CAG -3’ and AMP_R 5’-AAG GAG GTG ATC CAR CCG CA -3’ were used. PCR reaction mixture was prepared by mixing 25 µl of the Tag 2x Master mix (buffer, polymerase and dNTPs), forward primer 1 µl, reverse primer 1µl, and UPH_2_O 22 µl. Then, 49 µl of the mixtures were added to a sterile PCR tube; and 1 µl of the gDNA was used as template; the amplification reactions were carried out in a thermal cycler (Bio-Rad Mycycler).

#### DNA electrophoresis

PCR products were separated in a 1% agarose gel and stained with ethidium bromide followed by examination on UV illuminator and images were captured by a digital camera

#### Sequencing of the PCR products

A 16S rRNA PCR amplification and sequencing was performed by Eurofins, Novogene (Hong Kong). The V4 hypervariable region of the 16S rRNA was amplified using specific primers 515F and 806R. All the PCR reactions were carried out with Phusion® High-Fidelity PCR Master Mix (New England Biolabs). The libraries generated with TruSeq DNA PCR-Free Sample Preparation Kit were sequenced using paired-end Illumina sequencing (2 × 250 bp) on the HiSeq2500 platform (Illumina, USA).

#### Phylogenetic analysis

Forward and reverse sequences were assembled and edited using BioEdit Sequence Alignment Editor Version 5.0.9. Sequence similarity was estimated by searching the homology in the Genbank DNA database using BLAST. Finally, the isolate was then identified based upon the sequence.

The evolutionary history was inferred using the Neighbor-Joining method (27). The optimal tree with the sum of branch length = 0.19972900 is shown. The percentage of replicate trees in which the associated taxa clustered together in the bootstrap test (1000 replicates) are shown next to the branches. The tree is drawn to scale, with branch lengths in the same units as those of the evolutionary distances used to infer the phylogenetic tree. The evolutionary distances were computed using the Maximum Composite Likelihood method and are in the units of the number of base substitutions per site. All positions containing gaps and missing data were eliminated. There were a total of 891 positions in the final dataset. Evolutionary analyses were conducted in MEGA7.

##### Statistical analysis

All the measurements were performed in triplicate, and the results were expressed as mean standard deviation (SD). Data were analyzed by the one-way ANOVA plus post hoc Duncan’s test by SAS software (ver. 9.2, Raleigh, NC). The phylogenetic tree was prepared using MEGA7 (version 7.0) Statistical significance was determined at p < 0.05.

## Results

Isolation of potentially probiotic lactic acid bacteria from traditional fermented foods A total of 90 (30 from each sample) lactic acid bacteria were isolated from three different traditionally fermented Ethiopian food products (*Teff* dough, *Ergo* and *Kocho*). Among them, 56 (62.22%) isolates were found Gram-positive, endospore negative and catalase negative.

### Morphological, biochemical and physiological characterization

The 4 acid-bile-tolerant LAB isolates were identified through their morphological, biochemical and physiological features (S1Table). All the isolates were straight rod shaped, and able to grow at 4% NaCl salt concentration. From the 4 acid-bile-tolerant LAB isolates, 3 of the isolates were found growing at 6.5% NaCl salt concentration. Testing the abilities of isolates growing at 15 °C and 45°C indicated that 3 of the isolates were able to grow at both 15 °C and 45°C (S1Table). On the other hand, isolate E031 was not found growing at 45°C.

Out of the 4 acid-bile-tolerant isolates, 2 isolates (T035 and K011) produced gas from glucose (S1Table). Accordingly, the four isolates were found to be equally divided into homofermentative and heterofermentative types (50:50) (**Error! Reference source not found.**). Regarding arginine hydrolysis test, isolates E031, T035 and K011 were found positive for arginine hydrolysis, whereas E052 was not able to hydrolyze (**Error! Reference source not found.**).

#### *In vitro* characterization of probiotic properties

### Tolerance to low pH

Out of 56 isolates, 4 isolates (7.14%), 4 isolates (7.14%) and 9 isolates (16.07%) tolerated pH values of 2, 2.5 and 3 for 3 h, respectively (S2Table). Upon further extension of the incubation period to 6 h the 8 isolates (each 4 tested at pH 2.0 and 2.5) survived, whereas only 5 survived from 9 the extension of incubation period to 6 h at pH 3.0 (S2Table).

Therefore, out of the total 56 LAB isolates, 4 (7.14%) isolates were survived pH 2.0, 2.5 and 3.0 up on exposure for 3 and 6 hours and mean value of the treatments were significantly different at p<0.05 (Table). Among them, 1 (25%) was isolated from each *Teff* dough and *Kocho* sample. And also 2 (50%) LAB cultures were isolated from *Ergo* samples. In general, survival rate of the isolates was ranged from 38.40 to 97.11% at different pH values for 3 and 6 h incubation period (Table). Isolates, like E052, K011 and T035 were found highly tolerant and persisted above 50% for both 3 and 6 hours. On the other hand, isolate K031 was not able to grow above 50% at pH 2.0 up on exposure for 3 and 6 h (Table). Though the survival rate of the isolates was markedly reduced at pH 2.0 for 6 h exposure, all the isolates were taken to the next experiments.

#### Tolerance to bile salts

All of the 4 LAB isolates (E052, E031, K011 and T035) were able to survive above 90% in the presence of 0.3% of bile salt (S1Fig. *1*). Isolate E031 was the most tolerant with 97.22% survival rate followed by isolate E052 and T035 with 93.62% and 93.38% survival rate, respectively. However, isolate K031 was showed 91.37% survival rate (S1Fig. *1*).

### Antimicrobial activities

The diameter of inhibition zones showed that crude extracts from each isolate had antimicrobial effect against each tested foodborne pathogen (S4Table). The average zones of inhibition by which the crude extracts inhibited the growth of the test foodborne pathogens were found ranging between 17 to 21 mm. Isolate T035 displayed highest antagonistic activity against *Staphylococcus aureus, Listeria monocytogenes E. coli*, and *S. typhimurium* with inhibition zone ranged from 19.33 to 21 mm in diameters. Whereas, E052 showed minimum inhibition zone of diameter ranged from 17 to 19 mm against the indicator microorganisms (S4Table).

#### Antibiotic susceptibility test

The antibiotic susceptibility test of the selected LAB isolates to some common antibiotics showed sensitivity to tetracycline, ampicillin and erythromycin. However, all the 4 potential probiotic LAB isolates were displayed resistant to kanamycin and streptomycin (Table).

### Bacterial adhesion to stainless steel plates

The adherence ability of the potential probiotic LAB isolates found ranging between 33.48 and 36.30 % (S2 Fig. 2). Isolate K011showed the highest (36.30%) adherence rate followed by isolates T035 and E052 with 33.48% and 33.17% adherence rate, respectively. Whereas, isolate E031 showed the least (32.75%) adherence rate (S2 Fig. 2).

#### Identification of probiotic LAB isolates by 16S rRNA gene sequencing

The 16S rRNA gene sequences of the 4 LAB isolates with the best potential probiotic properties showed the highest homology to the known species of bacteria in the database (Fig.). Accordingly, E052 showed 99% match with *Lactobacillus plantarum* strain JCM 1149, T035 showed 99% homology with *Lactobacillus plantarum* strain CIP 103151, K011 showed 95% similarity with *Lactobacillus paracasei* subsp. tolerans strain NBRC 15906 and E031 showed 99% homology with *Lactobacillus paracasei* strain NBRC 15889 (Fig.).

## Discussion

The study of probiotic activities, for application in preservation of food products and human health remains to be of great interest. Currently, interest in antagonistic feature of probiotic LAB against foodborne pathogens has indicated their potential to be likely alternatives to chemical drugs (28). Therefore, a significant effort has been made to select lactic acid bacteria originating from the traditional Ethiopian fermented food products on the basis of the most important technological, functional and safety criteria in order to obtain potential probiotic lactic acid bacteria.

All the selected four potential probiotic bacteria were identified as LAB based on their morphological, biochemical and physiological characteristics that were found to be equally divided into homofermentative and heterofermentative types. This finding is in accordance with Mamo, Assefa (29) who isolated *Lactobacillus* species from *Ergo* and found that they were grouped as homofermentative and heterofermentative types. Akalu, Assefa (15) have also reported that the LAB isolated from *Shamita* and *Kocho* were identified morphologically, biochemically and physiologically and comprised of both heterofermentative and homofermentative types. Azadnia and Khan Nazer (30) reported that the lactic acid bacteria isolated from traditional drinking yoghurt in tribes of Fars province was able to grow at 4% NaCl concentration but not at 6.5% NaCl concentration.

In the present study, out of 56 probiotic LAB isolates tested to pH 2 for 3 and 6 h, only 4 (7.14%) isolates showed tolerance to pH 2 for 6 h (S1Table). Thus, the tolerance of the four *Lactobacillus* spp. to pH 2.0 for 6 h showed a survival rate of 38.40 to 73.29%. Similar to this study, Mourad and Nour-Eddine (31) have demonstrated that *Lactobacillus plantarum* OL12, *L. plantarum* OL9, *L. plantarum* OL15 and *L. plantarum* OL33 isolated from fermented olives showed survival percentage of 55%, 49%, 65% and 57%, respectively when exposed to pH 2.0 for 2. However, these results are not in accordance with those reported by Akalu, Assefa (15), Rajoka, Mehwish (32) who have indicated that most strains of *Lb. plantarum* isolated from different sources showed survival rate above 80% at pH 2 for 3h. Other reports revealed that 5 acid tolerant *Lactobacillus* strains showed above 89% survival rate after exposure to pH 2 for 3 h (33). Akalu, Assefa (15) verified that 14/17 *Lactobacillus* strains isolated from traditionally fermented *Shamita* and *Kocho*, were found tolerant to pH 2 when incubated for 6 h with survival rate of 74-89%. However, incubation at low pH resulted in significant decrease in the survival rate of all LAB isolates as noted in another study (Guo, Kim (34). The authors noted that the viable counts of all lactic acid bacteria were significantly affected by low acidity, especially at pH 2.

In this study, out of the 56 probiotic LAB isolates, only 4 (7.14%) isolates were also tolerated to pH 2.5 for 3 and 6 h (S1Table). Similarly, out of 56 LAB isolates, 9 isolates (16.1%) and 5 isolates (8.93%) were survived at pH 3 for 3 and 6 h, respectively. Thus, the survival rate of the four *Lactobacillus* strains at pH 2.5 and 3 for 3 and 6 h was found to be tolerant with different survival rate (77.98-97.11%) and (65.58-90.49%), respectively (S2Table). Similarly, Akalu, Assefa (15) have reported that out of 30 LAB isolated from traditionally fermented Ethiopian beverage and food (*Shamita* and *Kocho*), 17 lactobacilli showed 82 to 97% and 81 to 91% survival rate at pH 2.5 and 3 for 3 and 6 h, respectively. This is also similar to the report of Oh and Jung (33) who have showed a better survival rate (above 88%) of *Lactobacillus* species in pH 2.5 and 3 for 3 h that were isolated from Omegisool, a traditionally fermented millet alcoholic beverage of North Korea. As reported by Haghshenas, Nami (35), *Lactobacillus* strains isolated from Iranian fermented dairy products survived (71% to 76%) at pH 2.5 for 3 h. From the previous investigations, it has generally been accepted that an isolate with full tolerance to pH 3.0 for 3 h can be considered as high acid resistant strain with promising probiotic properties (34,36). Very recently, Tang, Qian (37) have reported that all the 9 *Lactobacillus plantarum* strains recovered from the feces of breast-feeding piglets were found to be highly tolerant to pH 3 for 3 h.

The present results are in contrast to Mamo, Assefa (29) who have found low to high survival rates (1.03–100%) for the six *Lactobacillus* species at pH 2.5 and 3.0 for 3 h. The same authors indicated that the maximum survival rate of the six strains were 22.5% at pH 3 for 6 h. Nevertheless, complete loss of viability of lactobacilli isolated from traditional Ethiopian *ergo* was recorded at pH 2.5 for 6 h (29). In addition, the current survival rate was higher than that of previously reported strains such *Lactobacillus plantarum* at pH 3 for 2 and 6 h exposure (31). The 4 acid-tolerant LAB isolates showed high tolerance to bile salt conditions (91.37% to 97.22%) (S1Fig. *1*). Bile salt tolerance is also considered as important selection criterion for probiotic isolates in order to survive the conditions in the small intestine. Moreover, tolerance to a high bile salt condition is also strain specific as demonstrated earlier where different *Lactobacillus* species isolated from Omegisool, a traditionally fermented millet alcoholic beverage in Korea showed considerable tolerance to bile salt (33). Similar to the present findings, the results in other studies have revealed that all the isolated strains displayed high tolerance to bile salt conditions and the survival rates of *Lactobacillus* strains ranged from 88% to 92% (35). In a related study, Akalu, Assefa (15) have also showed that out of the 30 tested LAB isolates, 17 *Lactobacillus* isolates obtained from Ethiopian traditionally fermented *Shamita* and *Kocho* showed remarkably high tolerance to an environment containing 0.3% bile salt. On the other hand, Boke *et al.* (2010) have indicated that *Lactobacillus* strains B3, G12, A13 and 22 exhibited low level of tolerance to 0.3% bile salts with survival rate of 36%, 33 %, 3% and 3%, respectively. In line with this, Handa (38) has reported that all of the LAB isolates demonstrated low level of tolerance to bile salts by displaying surviving percentage less than 50% when exposed to 0.3% bile salts after 24 h at 37°C.

The selected four potential probiotic lactic acid bacterial strains (*Lactobacillus plantarum* strain JCM 1149, *Lactobacillus plantarum* strain CIP 103151, *Lactobacillus paracasei subsp. tolerans* strain NBRC 15906 and *Lactobacillus paracasei* strain NBRC 15889) exhibited varying degree of antagonism against *Staphylococcus aureus, Listeria monocytogenes, Salmonella typimurium* and *Escherichia coli* (S4Table) According to Handa (2012), isolates having clearance zones ≤9 mm and ≥12 mm diameter against the test pathogens indicated poor and strong antimicrobial activity, respectively. Accordingly, all the selected potential probiotic LAB strains (n=4) exhibited strong antimicrobial activity against the foodborne pathogens, where T035 (*Lactobacillus plantarum* strain CIP 103151) displayed the highest antagonistic activity against *Staphylococcus aureus, Listeria monocytogenes, E. coli*, and *S. typhimurium* with the inhibition zone ranged from 19.33 to 21 mm in diameters.

In accordance to the current study, Bassyouni, Abdel-all (39) have demonstrated that all of the *Lactobacillus* isolates obtained from Egyptian dairy product had strong antibacterial effect against *E. coli* and *Salmonella typhimurium*. However, among the results that revealed by the same author, 3 isolates had the most potent antimicrobial activity against the tested pathogenic microorganisms with inhibition zone ranged from 17 to 21 mm in diameters. In agreement to this study, Tadesse, Ephraim (5) have verified that all the LAB isolates (n=118) originated from *Borde* and *Shamita* belonging to the genera *Lactobacillus, Lactococcus, Leuconostoc* and *Streptococcus* were found to inhibit the growth of the test strains such as *S. aureus, Salmonella* spp. and *E. coli* O157:H7 with inhibition zones that ranged from 15 to 17 mm in diameters. In line with this, Choi, Patra (40) have reported that out of the 4 strains of LAB, *Lactobacillus* strain was completely inhibited the growth of foodborne pathogens, *E. coli* O157:H7 ATCC 35150, *Salmonella enteritidis* KCCM 12021, *Salmonella typhimurium* KCTC 1925 and *S. aureus.* Tigu, Assefa (14) have also revealed that out of the 11 probiotic LAB isolated from traditional Ethiopian fermented condiments, namely *Datta* and *Awaze*, 2 *Lactobacillus* isolates inhibited the growth of *Salmonella typimurium* and *Escherichia coli* with inhibition zones ranging from 10.3 to 14.3 mm in diameters. In line with this, Haghshenas, Nami (35) have reported that among the selected 8 LAB isolated from Iranian fermented dairy products, *Lactobacillus* species, particularly *Lb. plantarum* 15HN, showed the most efficient antagonistic activity against *Staphylococcus aureus, Listeria monocytogenes, Salmonella typimurium* and *Escherichia coli* with inhibition zones of 11.7, 13.7, 12.3 and 12.3 mm diameters, respectively. Likewise, Rajoka, Mehwish (32) have verified that all the *Lactobacillus rhamnosus* isolated from human milk inhibited the growth of *Staphylococcus aureus, Salmonella typimurium* and *Escherichia coli* using agar well diffusion method with variable diameters (6 mm to 14 mm).

Disparity in the antagonism activity against different pathogens indicates that probiotic strains are highly pathogen specific and prerequisite for probiotic potential. In general, the antimicrobial activity of probiotics might be caused by the production of antimicrobial compounds such as organic acids, ethanol, carbon dioxide, hydrogen peroxide, short chain fatty acids, and bacteriocins. Therefore, by producing these antimicrobial compounds, probiotic microorganisms gain an advantage over other microorganisms to survive in the adverse conditions of gastrointestinal tract (38).

All of the tested four *Lactobacillus* strains were found to be resistant to streptomycin and kanamycin but sensitive towards tetracycline, ampicillin and erythromycin (Table). These results were found in agreement with the results that were obtained using *Lactobacillus* species (32). Tigu, Assefa (14) have also reported that all of the LAB isolates obtained from traditional fermented condiments such as *Datta* and *Awaze* were susceptible to ampicillin, erythromycin and tetracycline. Similarly, according to Amraii, Abtahi (41), all of the selected LAB isolates were sensitive towards ampicillin, erythromycin and tetracycline. On the contrary, Sukmarini, MUSTOPA (42) have reported that out of 120 isolates of LAB from four different Indonesian traditional fermented foods, 16 isolates were resistant to erythromycin. In line with this, Pan, Hu (43) observed that among the 12 *Lactobacillus* species obtained from Chinese fermented foods, 5 isolates were sensitive to kanamycin, 7 resistant to erythromycin, 9 resistant to ampicillin and 8 isolates were resistant to tetracycline.

Among the main important characteristics of probiotic bacteria, adhesion to the intestinal mucosa is required. The current results showed that the screened probiotic LAB isolates possessed *in vitro* adherence property to stainless steel plates with adhesion rate ranging from 32.75 to 36.30%. In agreement with the current study, El-Jeni, El Bour (25) have reported that the adhesion rate of lactic acid bacteria to stainless steel plates was ranged from 32 to 35%. Winkelströter, Gomes (44) have also revealed that pure culture of *Lb*. *sakei* ATCC 15521 showed strong adherence to stainless steel surface. Generally, this suggests that our potential probiotic LAB isolates may have a potential capacity to colonize the gastrointestinal (GI) tract mucosa.

The phylogenetic analysis and the 16S rDNA sequencing assigned all the LAB isolates with probiotic properties to genus *Lactobacillus* and the identified species were *Lactobacillus plantarum* strain JCM 1149, *Lactobacillus paracasei* strain NBRC 15889, *Lactobacillus plantarum* strain CIP 103151 and *Lactobacillus paracasei* subsp. tolerans strain NBRC 15906 (Fig.). According to Shokryazdan, Sieo (45), the results of comparative 16S rRNA gene analysis showed LAB isolates belonged to *L. acidophilus, L. fermentum, L. buchneri* and *L. casei*. In addition, Cho, Lee (46) have identified *Lactobacillus* strains with potential probiotic properties from the feces of breast-feeding piglets using 16S rRNA genes analysis. Similarly, other recent studies (Dowarah, Verma (6) have revealed the strain level identification of diverse lactic acid bacteria with potent probiotic properties isolated from some substrates using phylogenetic estimation of 16S rDNA genes.

## Conclusion

In the present study, four *Lactobacillus* strains isolated from *Ergo, Kocho* and *Teff* dough were found to have potentially probiotic characteristics. It is suggested that these strains can be a good candidate for food industries as prospective probiotic cultures with other human health benefits. However, further research work is needed to evaluate the *in vivo* probiotic characteristics of these potential lactic acid bacteria.

**Fig 1.**
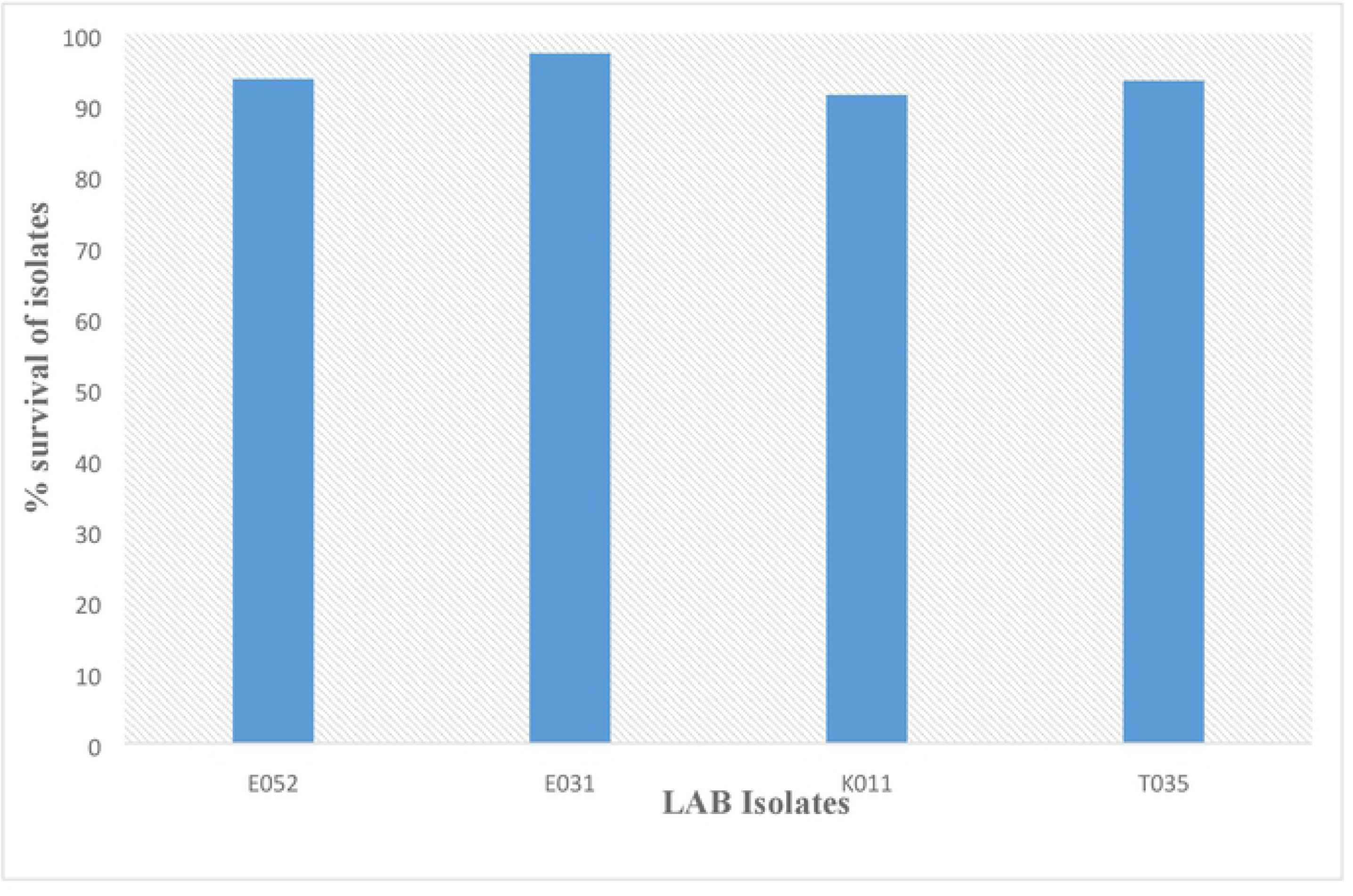
*Percentage survival of probiotic lactic acid bacteria against 0.3% bile salt. Means are significantly different at p<0.05. Results expressed as average (n* = *3)* ± SD (standard deviation).

**Fig 2.**
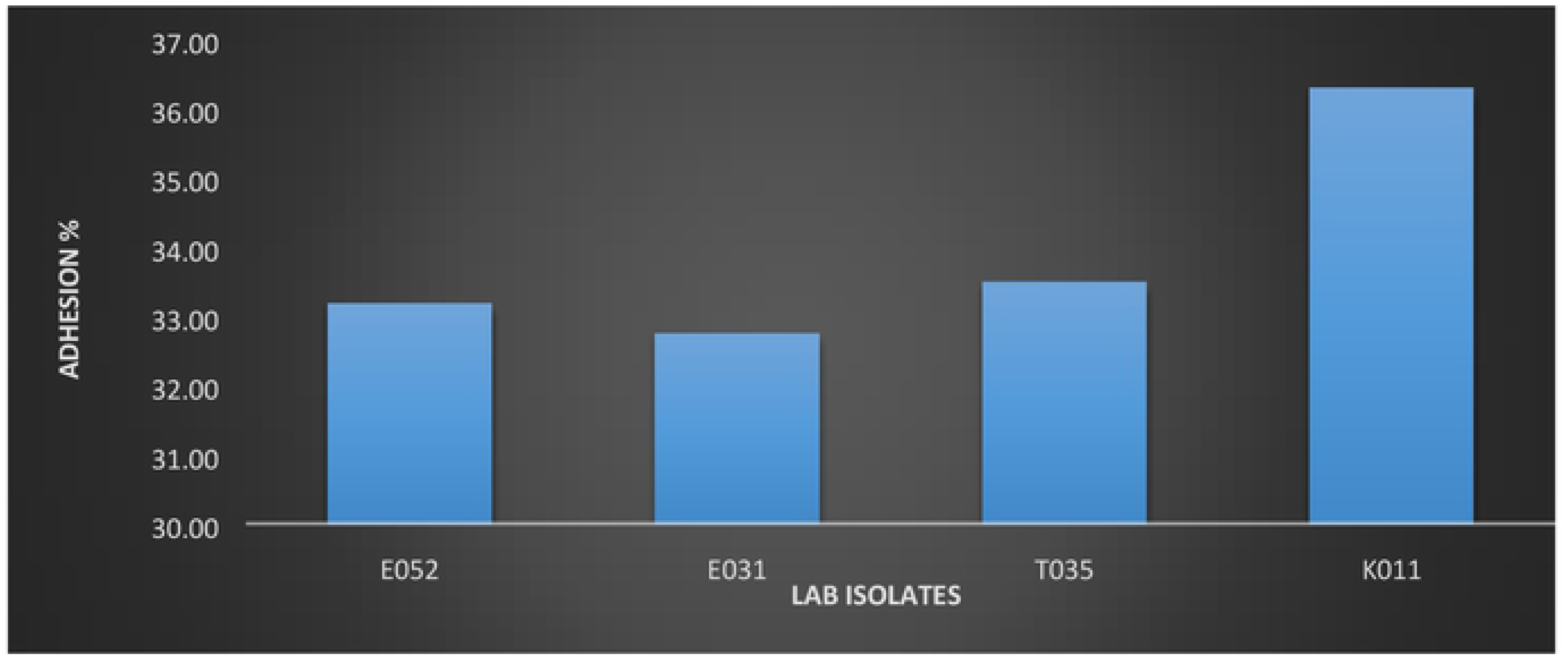
The adhesion of lactic acid bacteria to stainless steel plate. Means not connected by same letter are significantly different at *p*<*0.05.* Results expressed as average *(n =* 3) ± *SD (standard deviation).*

**Fig 3.**
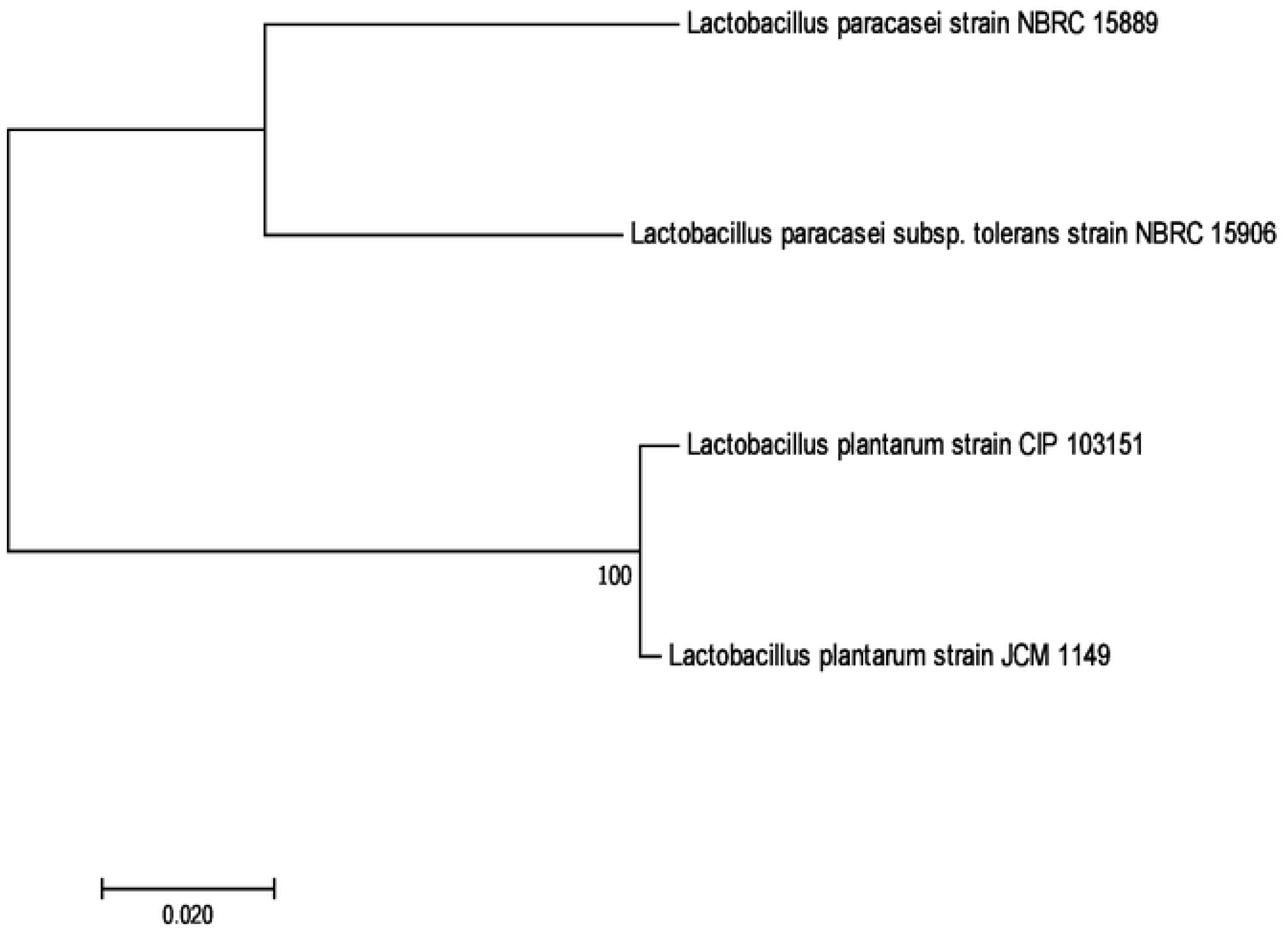
Evolutionary relationships of the isolation to the known strains.

## Acknowledgements

The authors acknowledge Microbial and Molecular Biology of Addis Ababa University and Helsinki University, Finland for provision of all laboratory facilities and Ethiopian Public Health Institute (EPHI), Addis Ababa, Ethiopia for providing test organisms.

## Supporting information

S1Table. Physiological, morphological and biochemical characteristics of the isolates.

S2Table. Acid tolerance patterns of probiotic LAB at different pH values after 3 and 6 h exposure.

*S3* Table. Percentage survival of probiotic lactic acid bacteria at different pH levels.

*S4Table. Antimicrobial activities of LAB against some foodborne pathogens*

S5 Table. Antibiotic susceptibility profile of probiotic LAB isolates.

*S1Fig. 1*. *Percentage survival of probiotic lactic acid bacteria against 0.3% bile salt.*

S2 Fig. 2. The adhesion of lactic acid bacteria to stainless steel plate.

*S3* Fig. Evolutionary relationships of the isolation to the known strains.

